# Individual Behavioral and Physiological Responses During Different Experimental Situations – Consistency over Time and Effects of Context

**DOI:** 10.1101/477638

**Authors:** Alexandra Safryghin, Denise V. Hebesberger, Claudia A.F. Wascher

## Abstract

In a number of species, consistent behavioral differences between individuals have been described in standardized tests, *e.g.* novel object exploration, open field test. Different behavioral expressions are reflective of different coping strategies of individuals in stressful situations. A causal link between behavioral responses and the activation of the physiological stress response is assumed but not thoroughly studied. Also, most standard paradigms investigating individual behavioral differences, are framed in a fearful context, therefore the present study aimed to add a test in a more positive context, the feeding context. We assessed individual differences in physiological (heart rate, HR) and behavioral responses (presence or absence of pawing, startle response, defecation, snorting) of twenty domestic horses (*Equus caballus*) in two behavioral experiments, a novel object presentation and a pre-feeding excitement test. Experiments were conducted twice, in summer and autumn. Both experiments caused higher mean HR in the first ten seconds after stimulus presentation compared to a control condition, but mean HR did not differ between the experimental conditions. Interestingly, in the novel object experiment, horses displaying stress-related behaviors during the experiments also showed a significantly higher HR increase compared to horses which did not display any stress-related behaviors, reflecting a correlation between behavioral and physiological responses to the novel object. On the contrary, in the pre-feeding experiments, horses that showed fewer behavioral responses had a greater HR increase, indicating the physiological response being due to emotional arousal and not behavioral activity. Moreover, HR response to experimental situations varied significantly between individuals, and although we found HR to be significantly repeatable across experiments, repeatability indices were low. In conclusion, our findings show that horses’ behavioral and physiological responses differed between test situations and that high emotional reactivity, shown via mean HR and HR increase, is not always displayed behaviorally.

## Introduction

The perception of a potential threat to homeostasis, caused by extrinsic or intrinsic stimuli (stressors) results in the activation of the physiological stress response in animals (Chrousus and Gold 1992; Moberg 1985). Regardless of the intensity of the stressor, individuals respond both behaviorally and physiologically, and these responses aim to counteract the effects of the stimuli and to re-establish homeostasis (Chrousus and Gold 1992). Physiological activation in response to a stressor causes a fast release of norepinephrine in the brain, triggering the activation of the sympatho-adreno-medullary (SAM) axis, and the hypothalamic-pituitary-adrenal (HPA) axis (Goldstein 2010). As a consequence, heart rate (HR) and glucocorticoid levels increase (Elia et al. 2010; McGreevy et al. 2005).

The way an individual perceives a stressor can significantly influence its responsiveness (Moberg 1985). Indeed, both endogenous and exogenous factors have been shown to modulate an animal’s sensitivity to a perceived threat, which results in individual differences in responsiveness within the same species (Gosling 2001). For example, the presence of constant environmental stressors can lead to a desensitization of the threat-perception, or learned helplessness, which can cause a reduction of individual responses to a stressor (Ellis et al. 2014). Furthermore, early differences in environmental stimuli, such as differences in maternal investment, may cause individuals to follow diverse trajectories, shaping their behavioral repertoire and physiological responsiveness in different manners (Claessens et al. 2011; Stamps 2003). For example, Meany (2001) showed how rat pups raised by low-licking mothers are more fearful and more sensitive to stress in adulthood compared to pups reared by high-licking mothers. Moreover, animals can be bred to either show high or low responsiveness to stressors, indicating that individual differences in stress-responsiveness are heritable (Carere et al. 2003; Flaherty and Rowan 1989). Studies on great tits (*Parus major*) have shown individual physiological stress responses to be related to differences in exploration strategies (Carere and van Oers 2004) and heritable throughout four generations (Drent et al. 2003). If individual differences in physiological and/or behavioral responses are found to be relatively stable across time and contexts, they are regarded as temperamental traits (Cockrem 2007; Goldsmith et al. 1987; Lansade et al. 2008; Le Scolan et al. 1997).

Temperamental traits have been found in both vertebrate and invertebrate taxa (Carere et al. 2003; Kralj-Fišer et al. 2010; Momozawa et al. 2003; Riechert and Hedrick 1993; Verbeek et al. 1994; While et al. 2009; Øverli et al. 2006). Studies have shown that individual differences along the shy-bold axis are linked with aggressiveness (Kralj-Fišer et al. 2010), exploratory behavior (Sibbald et al. 2009), neophobia (Momozawa et al. 2003), and physiological responsiveness, such as hormonal (Carere et al. 2003), and HR modulation (Kralj-Fišer et al. 2010). Heart rate presents a valid indicator of emotional arousal, defined as an internal state, which is triggered by specific extrinsic or intrinsic stimuli (Anderson and Adolphs 2014). Arousal can range between the subject being calm – low arousal, and excited– high arousal, as well as the experience being of positive or negative valence (Russell 1980). Briefer et al. (2015) have studied the effect of emotional arousal and valence on the physiological and behavioral response in goats, showing both, positive (feeding) as well as negative (frustration, isolation) emotional context to cause a significant physiological response compared to a control situation.

An evaluation of individual differences in emotional arousal in response to situations of positive emotional valence or the relationship of activation of the physiological stress response in situations of different emotional valence is still lacking. In the present study, we aim at investigating individual differences in emotional arousal in response to two experimental paradigms, a novel object exposure and a test of pre-feeding excitement. So far, most studies investigating temperamental traits in non-human animals have tested those in fearful contexts, *e.g.* novel object exploration or open field tasks (Dall and Griffith 2014; Smith and Blumstein 2008). Thereby we aimed to investigate physiological and behavioral responses in a fearful (novel object) and anticipatory (pre-feeding excitement) context.

In horses (*Equus caballus*), diverse factors influencing individual differences in behavior and physiology have been identified in terms of experience, such as habituation (Leiner and Fendt 2011), diet (Bulmer et al. 2015), handling (Visser et al. 2002), and maternal behavior (Houpt and Hintz 1983). Similarly, breed has also been found to strongly influence individual reactivity, suggesting a relationship between individual responsiveness and the heritability of traits (Hausberger et al. 2004; Lloyd et al. 2008). Further studies on equine temperament have focused on the assessment of horse responsiveness to different stimuli such as diverse environmental conditions (McCall et al. 2006; Schmidt et al. 2010a; Schmidt et al. 2010b), novel situations (Ellis et al. 2014; Fureix et al. 2009; Leiner and Fendt 2011; Visser et al. 2001; Visser et al. 2002) human interactions in terms of both handling (Ellis et al. 2014; Fureix et al. 2009; König von Brostel et al. 2011) and riding (Visser et al. 2008) and have shown how an individual’s response to threat – or ‘fearfulness’ – is stable across time (Lansade et al. 2008; Visser et al. 2001; Visser et al. 2003). However, contrasting results have been found according to the relationship between HR and behavioral parameters (Christensen et al. 2005; Lansade et al. 2008; Momozawa et al. 2003). For instance, only Momozawa et al. (2003) found behavioral correlations with HR in their fear-inducing experiments; results which were not supported in subsequent research (Christensen et al. 2005; Lansade et al. 2008).

In the present study, we aim at gaining further understanding of the relationship between individual differences in behavioral and physiological reactivity in horses. Heart rate represents a standardized, objective and non-invasive measure to infer an individual’s emotional arousal level, readily available to inexperienced behavioral observers (Lansade et al. 2008; Visser et al. 2003). We question whether physiological responses during a novel object presentation and pre-feeding test are caused by behavioral changes or purely emotional arousal, in the absence of behavioral activity. Furthermore, we ask whether behavioral and physiological responses are stable across time and contexts. We expect that horses show a greater physiological reaction to a fear-inducing situation such as being exposed to a novel object, compared to the anticipatory pre-feeding exposure.

## Methods

### Animals and housing

The study was conducted at the equine yard of the College of West Anglia (United Kingdom) between July and November 2017. The research was conducted on 20 horses which were individually stabled in loose boxes. Five of the 20 horses were tested only in one of the two experimental conditions, two solely for pre-feeding excitement and three only for novel object test, due to their lack of availability during testing periods (Table 1). The sample included 14 geldings (Age: mean 11.8 ± 3.8 years (yrs), range 6 - 18 yrs) and 6 mares (Age: mean 11.8 ± 2.6 yrs, range 10-17 yrs) of diverse breeds, use, and training experiences (Table 1). The horses were fed twice a day: once in the morning (0800-0830) and once in the afternoon (1500-1600). Water was available *ad libitum*, and feces were removed from the stables after the horses were fed. In the late afternoon (1600-1700), some of the horses were turned out in the paddock for the night.

**Table 1.** Age in years, sex and breed of the 20 horses tested for this study. Mean heart rate and standard deviation during the different experimental situations as well as whether the horses were categorized as high (orange) or low (blue) behavioral responders.

| ID | Age | Sex | Breed | NO 1 | NO2 | PF1 | PF2 |
| --- | --- | --- | --- | --- | --- | --- | --- |
| 1 | 13 | Gelding | Cob | NA | NA | 48.29 ± 9.89 | 57.14 ± 12.43 |
| 2 | 14 | Gelding | Welsh Pony | 51.47 ± 8.87 | 51.21 ± 5.66 | 67.27 ± 12.41 | 54.42 ± 8.01 |
| 3 | 17 | Mare | Gypsy Cob | 54.77 ± 9.71 | 45.16 ± 3.44 | NA | NA |
| 4 | 8 | Gelding | Cob | 44.54 ± 1.56 | 38.37 ± 0.40 | 41.08 ± 5.62 | 33.69 ± 2.48 |
| 5 | 14 | Gelding | Cob | 46.82 ± 2.69 | 54.67 ± 3.85 | 51.29 ± 9.89 | 55.47 ± 10.65 |
| 6 | 9 | Gelding | KWPN | 43.87 ± 6.93 | 45.93 ± 12.30 | 39.01 ± 2.80 | 39.95 ± 7.07 |
| 7 | 10 | Mare | Welsh Crossbred | 62.40 ± 12.40 | 43.16 ± 1.92 | 48.06 ± 5.75 | 65.76 ± 14.01 |
| 8 | 16 | Gelding | Welsh Crossbred | 71.47 ± 18.82 | 44.41 ± 2.40 | 68.63 ± 9.19 | 53.98 ± 13.29 |
| 9 | 13 | Gelding | Cob | NA | NA | 55.57 ± 5.06 | 50.73 ± 1.53 |
| 10 | 12 | Gelding | KWPN | 54.30 ± 5.87 | 40.24 ± 8.05 | NA | NA |
| 11 | 16 | Gelding | Shire X Warmblood | 83.26 ± 20.00 | 80.03 ± 29.02 | 60.78 ± 7.80 | 46.88 ± 6.05 |
| 12 | 11 | Mare | Appaloosa X Cob | 42.84 ± 1.54 | 41.92 ± 8.30 | 60.89 ± 3.36 | 48.30 ± 12.39 |
| 13 | 11 | Gelding | Welsh Section A | 62.73 ± 20.70 | 55.29 ± 25.91 | NA | NA |
| 14 | 12 | Mare | Thoroughbred X Cob | 43.93 ± 3.47 | 47.27 ± 4.77 | 61.14 ± 7.16 | 46.12 ± 3.40 |
| 15 | 18 | Gelding | Thoroughbred | 44.43 ± 8.47 | 36.94 ± 4.96 | 41.58 ± 1.78 | 33.59 ± 1.90 |
| 16 | 10 | Mare | Cob | 44.28 ± 12.98 | 66.93 ± 15.66 | 42.84 ± 4.98 | 44.89 ± 7.01 |
| 17 | 6 | Gelding | KWPN | 36.67 ± 2.59 | 31.41 ± 1.52 | 42.10 ± 4.30 | 38.41 ± 2.57 |
| 18 | 9 | Gelding | Irish Sports Pony | 69.63 ± 15.71 | 62.12 ± 8.10 | 64.30 ± 12.74 | 36.77 ± 3.04 |
| 19 | 11 | Mare | Warmblood | 40.25 ± 4.11 | 42.52 ± 6.27 | 50.69 ± 17.53 | 38.71 ± 4.06 |
| 20 | 6 | Gelding | Cob | 44.22 ± 9.89 | 39.63 ± 4.86 | 53.74 ± 8.18 | 37.48 ± 3.31 |

### Experimental design

We conducted two experimental tests – the pre-feeding excitement and the novel object test, presented in random order, with one trial per horse and repeated in summer (July/August) and in autumn (October/November). Behavior was recorded by video camera (Canon Legira HF R56), and HR was recorded using a Polar V800 system belt, placed around the chest of the horses. The belt consists of an electrode belt with a built-in transmitter, connected via Bluetooth to a wristwatch (receiver). To optimize the contact between the belt and the skin, both the coat of the horse in the interested area and the belt were wetted. The receiver was placed at the stable entrance or inside the stable. Prior to each test, an adjustment period of 5 minutes was allowed to let the horse habituate to the belt.

The pre-feeding excitement test was conducted during morning feeds (0800-0830) and started with the horses being shown a bucket containing their individual mix of hard feed on the floor outside the stable, while the other horses were fed. The horses’ physiological and behavioral responses were recorded for five minutes. Thereafter, the horses were fed by placing the feeding bucket inside the box and their behavioral and physiological responses were measured for the following ten minutes.

In the novel object test, the horses were exposed to two of three different objects in their stable. The first object was formed of a main cylindrical hard body (approximately 30 cm in length) filled with gravel which was fixed to a soft foam rubber ball (about 15 cm in diameter) and covered in fabric. The second object was formed of two cylindrical plastic tubes fixed together to form an ‘x’. Similar to the first object, the cylinders were approximately 30cm in length, filled with gravel and covered with fabric. In addition, twelve tennis balls of different colors and materials were pierced and attached to 4 strings (3 balls per string) of approximately 50 cm in length. These were then tied to the main body of the object and left hanging. The third object was an inflatable guitar of approximately one meter in length. All objects were attached to a string, around four-meters-long, to allow their retrieval from the stable.

The object assigned to each individual was randomized for each season as well as the order of the horses tested. To avoid the horses seeing the object before testing, the objects were covered from sight when carried around the yard. The novel object tests took place between the hours of 0900 and 1300 and between 1500 and 1800 when the yard was quiet, and the horses were fed. The testing procedure was based on the one used by Dai et al. (2015) and Górecka-Bruzda et al. (2011) and adapted for the present experiment. A novel object was placed over the box entrance, with the cord hanging over the stable door to keep the object at the height of approximately one meter. The object was kept in this position for the following five minutes and was then dropped to the floor (the objects filled with gravel created a muffled noise). The horse reaction was recorded for the following five minutes. Thereafter, the object was removed from the stable, whilst behavioral monitoring and HR measurement continued for another 15 minutes.

### Ethical statement

All applied methods were non-invasive, and the experimental procedure was approved by Anglia Ruskin University’s Departmental Research Ethics Panel.

### Data processing

Raw HR data were purged with a moving average filter to remove biologically implausible outlier values. The following HR variables were calculated: (1) mean HR in beats per minute (bpm) for the ten seconds preceding and following the food presentation, as well as preceding (control) and following the presentation, drop and removal of the object; (2) HR increase in bpm following the food presentation and novel object presentation, drop, and removal, calculated as difference between maximum value and three seconds average HR before the presentation of the stimuli.

Behavioral responses of the horses were analyzed from videos using SOLOMON Coder v. beta 17.03.22 (András Péter, www.solomoncoder.com). The behavior of the individuals was analyzed for the five minutes prior to the presentation of the hard feed for the pre-feeding excitement. For the novel object task, the five minutes following the presentation and drop of the object and the two minutes following its removal were analyzed. Behavior was recorded as continuous variables, *e.g.*, feeding as duration of behavior in s per observation period, or frequency of behavior per observation period, *e.g.*, snorting. The classification of the individual in high and low behavioral respondents was based on the frequency and duration of vocalizations, pawing behavior, startle response and defecation that the individuals performed during the experimental settings (Table 2).

**Table 2.**
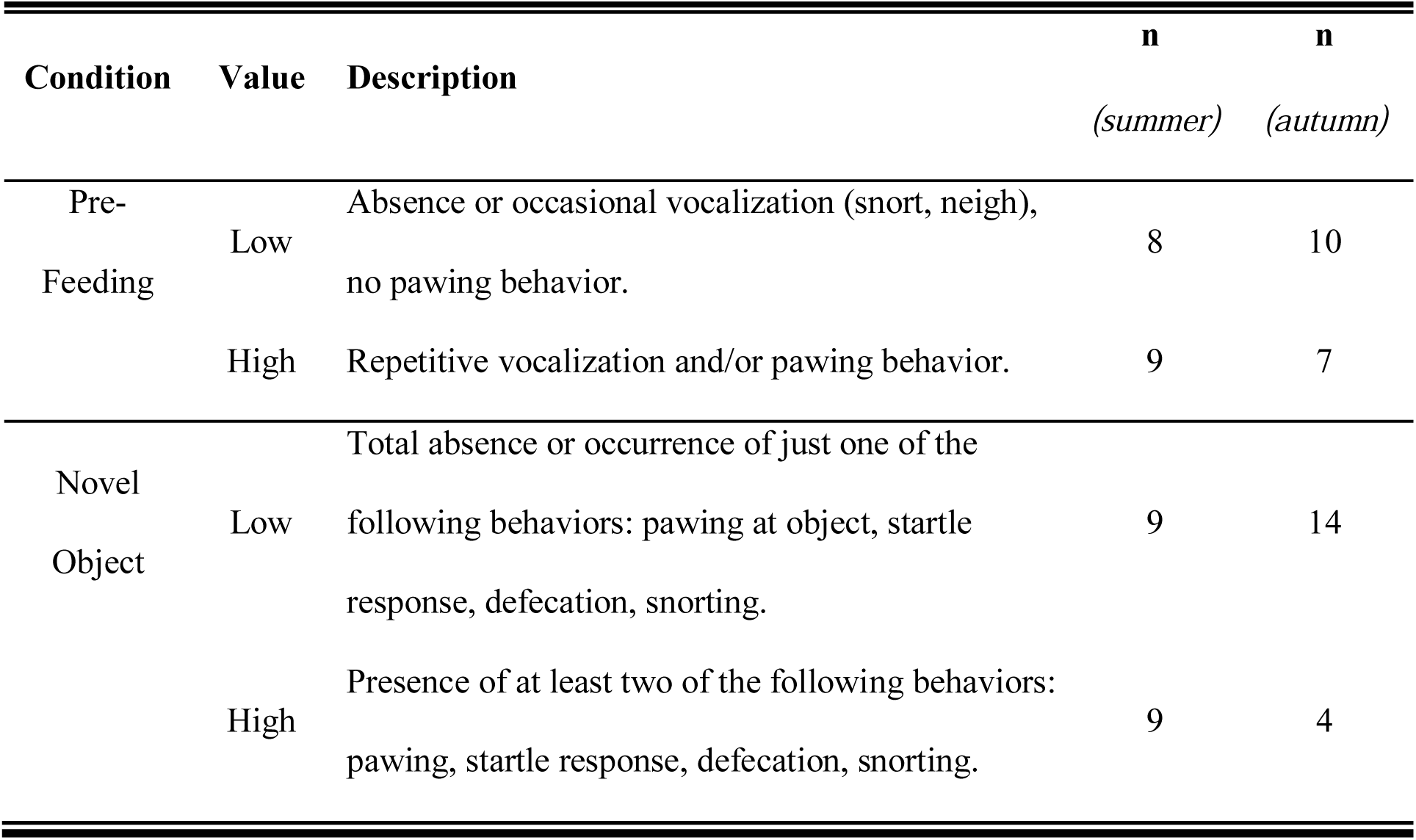
Description of behavioral categories and number of individuals per category, per season.

### Statistical Analysis

All data were analyzed using R version 3.4.3 (R Core Team, 2017; RStudio Team 2016). Generalized linear mixed models (GLMM) were conducted with the additional packages ‘glmmADMB’ (Skaug et al. 2016). For the comparison of HR, two generalized linear models were calculated. Experiment (pre-feeding excitement versus novel object), test period and behavioral category (high versus low respondents) were considered as fixed factors in both models, together with the interaction between behavioral category and experimental condition. For the purpose of the analysis, the different conditions of each experiment were individually included in the dataset, resulting in horses having multiple values per each experimental condition. The response variable was assigned to the ten second average HR (GLMM1) and the HR increase (GLMM2). Tukey test for multiple comparisons was chosen to gain further understanding of the effect of the categories of the fixed factors in the models. Specifically, the ‘multicomp’ (Hothorn et al. 2008) package was used to conduct the post hoc analysis. Moreover, we used a likelihood ratio test to compare models fit according to presence or absence of the individual random effect. Also, individual repeatability of the physiological and behavioral measurements was assessed using the ‘rptR’ package (Stoffel et al. 2017). In particular, we assessed the repeatability of the ten second average HR and HR increase with 1000 permutations for the physiological reactivity data collected for the control, novel object and pre-feeding conditions. The significance level was set at α = 0.05.

## Results

### Seasonal variation and behavioral categorization

Horses had a significantly higher mean HR during summer compared to autumn (GLMM1: z = 2.713, p = 0.007), but HR increase was not significantly different between the seasons (GLMM2: z = −0.94, p = 0.348). Out of the 20 horses tested, only four individuals were consistently categorized as high behavioral responders and one individual as low behavioral responder, in both experiments (novel object, pre-feeding excitement) and seasons (summer and autumn; see Table 1 for details).

### Physiological and behavioral responses to experimental situations

Average HR of the horses was significantly higher during the novel object experiment compared to the control period (Tukey: z = 5.205, p < 0.001; Figure 1A, Figure 2) and tended to be higher during the pre-feeding excitement compared to the control period (Tukey: z = 2.197, p = 0.0662; Figure 1A, Figure 2). Heart rate between novel object and pre-feeding excitement was not significantly different (Tukey: mean: z=-1.887, p=0.133; Figure 1A, Figure2), despite horses showing a significantly lower HR increase in the pre-feeding condition (GLMM2: z= −3.59, p < 0.001; Figure 1B). We found a significant interaction between behavior and experiment affecting HR. Mean HR during the novel object experiment was significantly higher in the group of horses showing a high behavioral response compared to horses showing a low behavioral response (GLMM1: z = −3.139, p = 0.002; Figure 1A). This pattern reversed regarding HR increase in the pre-feeding experiment, with the horses showing a low behavioral response having a higher HR increased compared to individuals with a high behavioral response (GLMM2: z = 3.56, p < 0.001; Figure 1B). Finally, the models including individual identity as random factor had a significantly better fit compared to the models without the random effect (ANOVA: mean HR: Deviance = 14.632, df = 1, p = 0.001; HR increase: Deviance = 7.984, df = 1, p = 0.005), indicating that the random effect also significantly variance in the data.

**Figure 1.**
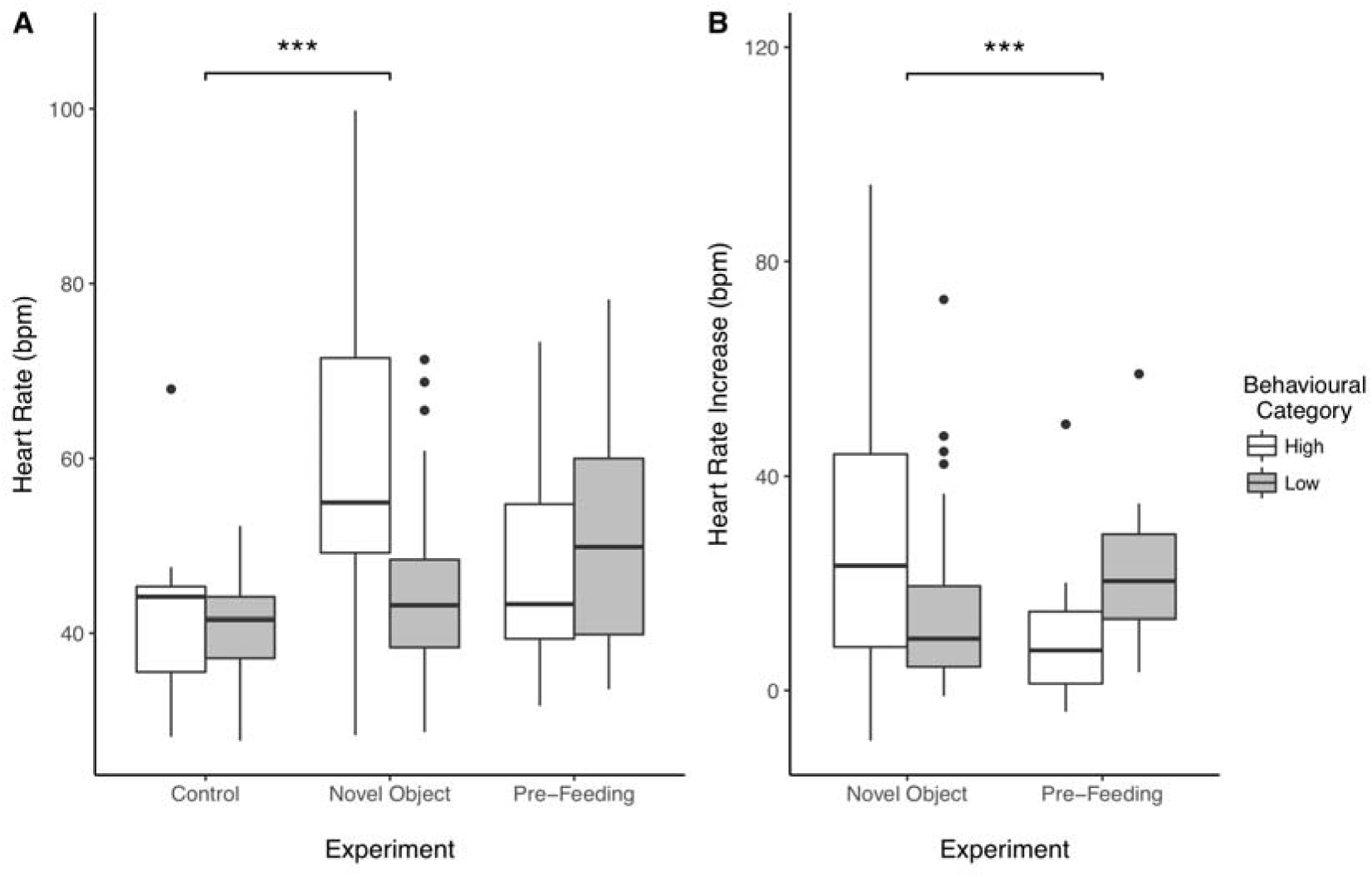
Effect of behavioral categorization on the mean heart rate (HR) (A) and HR increase (B) of the horses recorded during the study. Boxplots represent the median (black bar), the interquartile range – IQR (boxes), maximum and minimum values excluding outliers (whiskers) and outliers (black dots).

**Figure 2.**
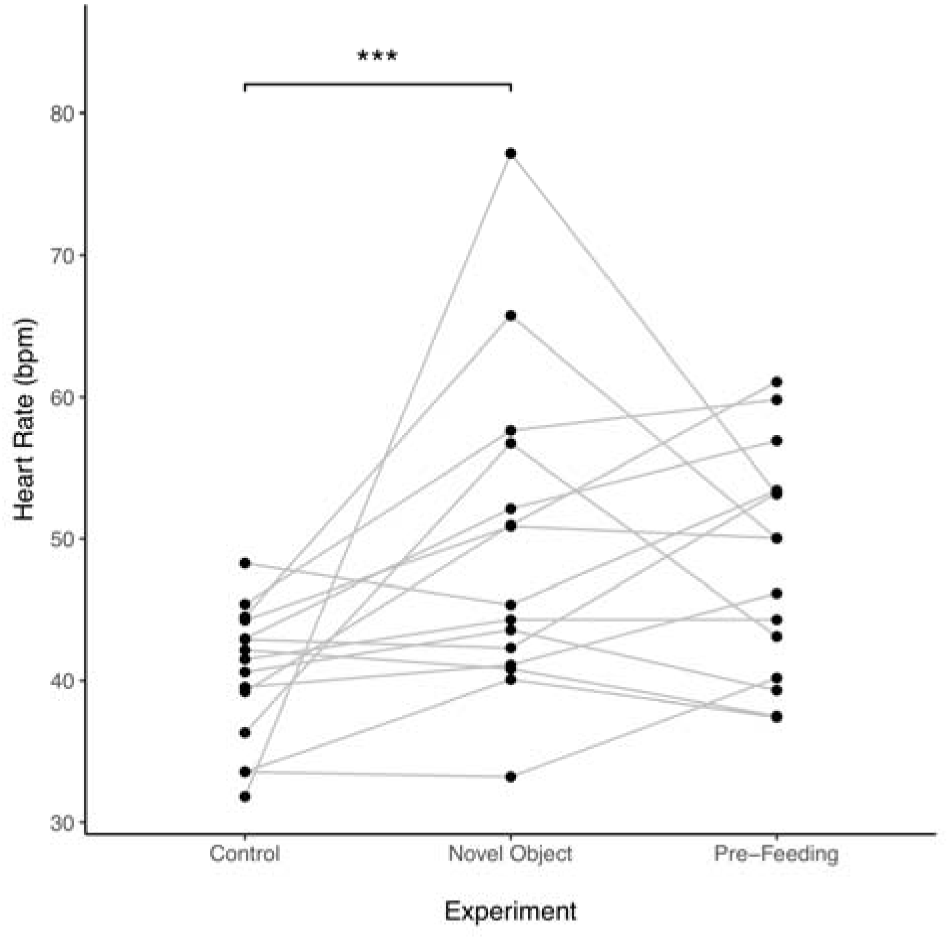
Individual average heart rate of horses across experiments. Points represent a value per individual horse and the lines connect individual horses over different experiments.

### Repeatability

Individual horses’ HR was significantly repeatable across experiments. However, the repeatability indices were low for both parameters tested, with the highest repeatability score recorded for the HR increase (*R* = 0.265, CI 95% [0.07, 0.442], p < 0.001); followed by mean HR (*R* = 0.201, CI 95% [0.06, 0.343], p < 0.001).

## Discussion

In the present study, we investigated individual behavioral and physiological responses of horses during two experimental procedures, a novel object experiment and a pre-feeding excitement test. Surprisingly, we found rather little behavioral consistency across tests and only a limited number of individuals responded similarly across contexts and seasons, which would have been expected when behavioral responses in experimental tests are indicative of temperamental traits. In contrast to behavior, heart rate (HR) was significantly repeatable across experiments, although it has to be pointed out that the repeatability indices were low. This indicates that physiological responses to experiments were consistent over time and contexts, as expected for temperamental traits. Repeatable individual variation in behavior, *i.e.* personality has been very much in focus of scientific research in recent years and was described in a vast variety of species, including horses (Grajfoner et al. 2010; Momozawa et al. 2005), however most of these studies are focusing on behavior rather than underlying physiological processes. For example, Grajfoner et al. (2010) compared individual horse behavioral ratings between high and low performers, showing how the combination of multiple traits, such as a horse being “nice”, “patient”, “easy to handle”, shape the perceived personality of horses. On the contrary, only a few studies investigated individual differences in physiological responses and their relationships with behavioral reactivity in non-human animals (Briefer et al. 2015; Ellis et al. 2014).

We described context dependent links between behavioral and physiological response. The two conducted experiments differed regarding their context, with the novel object experiment being considered as a fear-inducing situation (Dalmau et al. 2009; Lansade et al. 2008; Leiner and Fendt 2011), whereas the pre-feeding excitement test presents an anticipatory context (Mahnhardt et al. 2014). In the novel object experiment, a classic temperament test in non-human animals’ personality research (Carter et al. 2013), individuals who were behaviorally active also showed a higher average HR. This is especially relevant as the presentation of a novel object can be seen as a fear-inducing situation (Leiner and Fendt 2011). The results of the present study confirm that increased physical activity resulting from the performance of stress-related behaviors (*e.g*., startle response) correlates with higher mean HR compared to individuals not responding behaviourally in the experiment. It is difficult to conclude whether the physiological activation is caused by increased emotional arousal, *i.e.* the perception of fear or the increased physical activity, *e.g.* locomotion (von Borell et al. 2007; Visser et al. 2002).

A different pattern became apparent during the pre-feeding excitement task. Here, individuals lacking a behavioral response during the experiment showed a higher HR increase. This indicates that emotional arousal, but not physical activity, accounted for the increase in HR. Horses that were classified as highly behaviorally responsive during the pre-feeding experiments had an undetectable increase in HR, reflecting that physical activity may have increased their HRs before the presentation of the feed. Once the feed was presented their arousal level may not have changed in amplitude, as no significant HR increase was recorded, but possibly in valence, from negative to positive. Effects of emotional arousal on physiological responses have already been identified in other non-human animals. For example, in the study by Wascher et al. (2008), immobile greylag geese (*Anser anser*) watching aggressive interaction between conspecifics showed a significantly higher increase in HR compared to geese watching non-social interactions. Moreover, an increase in physiological reactivity resulting solely from emotional arousal was also identified in guide dogs (Fallani et al. 2007).

Our findings show that HR in the control situations did not necessarily predict HR during the experiments and horses differed in the strength of their responses, with some even decreasing HR. Such variation, together with the repeatability of the physiological reactivity, reflected how behavioral responses of horses do not necessarily predict physiological reactions during a novel object and pre-feeding excitement test. This is in line with previous studies on horses describing that individual classified as calmer had higher HR compared to more excited individuals despite showing less behavioral signs of stress when tested for pre-feeding reactivity (Ellis et al. 2014), as well as with other studies on cattle (*e.g.*, Christensen et al. 2005; Jezierski et al. 1999; Lansade et al. 2008; Welp et al. 2004).

Classical models regarding individual differences in behavior and physiology, assume them to be associated with each other to form different coping styles (Koolhaas et al. 1999). However, evidence for independent modulation of the hypothalamic-pituitary-adrenal axis (Boulton et al. 2015; Dosmann et al. 2015; Ferrari et al. 2013), the sympatho-adreno-medullary axis (Qu et al. 2018), and behavioral traits have been recently accumulating. Our study provides further evidence that HR and behavior might be regulated independently.

To conclude, our study illustrates the importance of accounting for context when studying individual differences in non-human animals, and we suggest that the inclusion of tests in non-fear related contexts is desirable. Further, we point at the importance of studying interconnectedness and independencies between physiological and behavioral reactions.

## Acknowledgements

We would like to thank Elizabeth Ragg and Nikki Bentley from the College of West Anglia for granting us access to their facilities and horses to conduct our experiments. Furthermore, we would like to thank all members of staff for helping us to facilitate our research on site. Finally, we would also like to thank Irene Susini for discussion and comments on the manuscript.

## Author Contribution statement

A.S., D.V.H. & C.A.F.W. designed the experiments

A.S. & D.V.H. conducted the experiments

A.S. & C.A.F.W. analyzed the data

A.S., D.V.H. & C.A.F.W. wrote the paper

